# The ketogenic diet is not effective in preclinical models of IDH1 wild-type and IDH1 mutant glioma

**DOI:** 10.1101/2020.06.23.166652

**Authors:** Rodrigo Javier, Craig Horbinski

## Abstract

Infiltrative gliomas are the most common neoplasms arising in the brain, and remain largely incurable despite decades of research. A subset of these gliomas contains mutations in *isocitrate dehydrogenase 1* (IDH1^mut^). This mutation disrupts cellular biochemistry, and IDH1^mut^ gliomas are generally less aggressive than IDH1 wild-type (IDH1^wt^) gliomas. Some preclinical studies and clinical trials have suggested that a ketogenic diet (KD), characterized by low-carbohydrate and high-fat content, may be beneficial in slowing glioma progression. However, not all studies have shown promising results, and to date, no study has addressed whether IDH1^mut^ gliomas might be more sensitive to KD. The aim of the current study was to compare the effects of KD in preclinical models of IDH1^wt^ versus IDH1^mut^ gliomas. *In vitro*, simulating KD by treatment with the ketone body β-hydroxybutyrate had no effect on the proliferation of patient-derived IDH1^wt^ or IDH1^mut^ glioma cells. Likewise, KD had no effect on the *in vivo* growth of these patient-derived gliomas. Furthermore, mice engrafted with Sleeping-Beauty transposase-engineered IDH1^wt^ and IDH1^mut^ glioma showed no difference in survival while on KD. These data suggest that IDH1^mut^ gliomas are not more responsive to KD, and that clinical trials further exploring KD in this subset of glioma patients may not be warranted.

## INTRODUCTION

Diffusely infiltrative gliomas strike over 17,000 people in the United States per year [1]. The vast majority of these tumors recur and progress, despite advancements in surgery, chemotherapy, radiotherapy, and immunotherapy. In 20-39 year-olds, gliomas are the 2^nd^ most common cause of cancer death in men, and are the 5^th^ most common cause in women [1]. The most common type of primary brain cancer in adults is diffusely infiltrative glioma; the most common subtype of infiltrative glioma, glioblastoma (GBM), is unfortunately also the most lethal. Despite great advances in treating many other kinds of cancer, the median survival of GBM patients is still only 12-15 months after diagnosis, even with surgical resection, radiation, and temozolomide therapy [2, 3]. Long-term prognosis is grim; only about 15% of patients with an infiltrative glioma survive 5 years after diagnosis. As a group, primary brain cancers rank #1 among all cancers in terms of average years of life lost [4]. Despite intensive research, very little progress has been made in the treatment of GBM, and new approaches are badly needed.

Alterations in cell metabolism have long been known to be a hallmark of cancer, ever since Otto Warburg first described the preferential reliance of cancer cells on aerobic glycolysis over oxidative phosphorylation [5, 6]. However, it was only relatively recently that mutations in metabolic genes were found to occur in some cancers. For example, approximately 20-30% of infiltrative gliomas carry mutations in isocitrate dehydrogenase 1 (IDH1) or, far less commonly, IDH2 [7-9]. This subset of gliomas tends to occur in grade 2-3 gliomas, disproportionately arises in younger adults, and is associated with longer survival. Wild-type IDH1 and IDH2 encode enzymes that catalyze the oxidative decarboxylation of isocitrate to α-ketoglutarate in the cytosol/peroxisomes and mitochondrion, respectively. In the process, these enzymes also generate reduced nicotinamide adenine dinucleotide phosphate (NADPH). Point mutations in codon 132 of IDH1 (usually R132H), and codon 172 of IDH2, cause the mutant enzymes to instead reduce α-ketoglutarate to D-2-hydroxyglutarate (D2HG), thereby consuming NADPH [10].

Because cancers mostly rely on glucose for energy and anabolism, numerous studies have explored the therapeutic potential of dietary carbohydrate restriction in cancer patients. This is achieved through a prolonged ketogenic diet (KD), which is characterized by high-fat, low-carbohydrate, and adequate-protein content. KD limits the bioavailability of carbohydrates and induces the liver to produce ketone bodies, such as β-hydroxybutyrate and acetoacetate, which are then converted into acetyl-CoA for use in the tricarboxylic acid (TCA) cycle [11]. The goal of KD is to force tumor cells to use ketone bodies instead of glucose, while still meeting the patient’s basic nutritional needs.

KD has been tested as an adjuvant therapeutic strategy in a number of cancers, including GBM, with mixed results thus far [12]. However, an aspect of this research that has not yet been addressed is whether patients with IDH1^mut^ gliomas might be particularly responsive to KD. One study suggested that the D2HG product of IDH1^mut^ can actually bind and inhibit ATP synthase, thereby inhibiting oxidative phosphorylation and ATP production [13]. In that study, human colorectal HCT116 cells artificially overexpressing IDH1 R132H were highly vulnerable to glucose deprivation *in vitro*, and were not able to use ketone bodies as effectively as IDH1^wt^ HCT116 cells. Therefore, we sought to explore whether KD might preferentially inhibit the growth of IDH1^mut^ gliomas *in vitro* and *in vivo*, using both patient-derived endogenous IDH1^wt^ and IDH1^mut^ xenografts in immunocompromised mice, as well as a Sleeping Beauty transposase-engineered model of IDH1^wt^ and IDH1^mut^ gliomas in immunocompetent mice.

## METHODS

### Ethics Statement

This study was performed in accordance with the recommendations in the Guide for the Care and Use of Laboratory Animals of the National Institute of Health. The protocol was approved by the Institutional Animal Care and Use Committee of Northwestern University (protocol #5715). All surgery was performed under isoflurane inhalant anesthesia. Every effort was made to minimize animal suffering.

### Cell Lines and Cell Culture

Five GBM cell types were derived from Dr. Jann Sarkaria at the Mayo Clinic Brain Tumor Patient-Derived Xenograft National Resource (https://www.mayo.edu/research/labs/translational-neuro-oncology/mayo-clinic-brain-tumor-patient-derived-xenograft-national-resource). Three were IDH1^wt^ (GBM6, GBM12, and GBM43), and 2 were IDH1^mut^ (GBM164 and GBM 196). Two additional IDH1^mut^ cell types were TB09, an anaplastic astrocytoma obtained from Dr Hai Yan at Duke University, and HT1080, a fibrosarcoma cell line from the American Type Culture Collection (ATCC). All IDH1^mut^ cells were R132H except for HT1080, which was R132C IDH1. NPA and NPAC1 were rodent isogenic cell lines engineered using the Sleeping Beauty transposon system, gifted courtesy of Dr. Maria Castro from the University of Michigan. Both NPA and NPAC1 have activating mutations in *NRAS* and inactivating mutations in *TP53* and *ATRX*; NPAC1 also expresses IDH1 R132H [14]. All IDH1^mut^ cell types produced high amounts of D2HG via liquid chromatography-mass spectrometry, relative to the IDH1^wt^ cells (not shown). All cell types are authenticated annually via short tandem repeat analysis.

GBM6, GBM43, TB09, and HT1080 cells were grown in Dulbecco’s modified eagle medium (DMEM, Corning) supplemented with 10% fetal bovine serum and 1% penicillin streptomycin at 37°C with 5% CO2.

### In vitro proliferation

Cells were plated in 24 well plates at a concentration of 5×10^4^ cells per well in triplicate. Experimental wells were supplemented with 10 mM β-hydroxybutyrate (Sigma Product #H6501). To mimic the low glucose environment characteristic of physiological ketosis, a formulation of DMEM (Corning) with 1.0 g/L of glucose was tested alongside the normal glucose concentration of 4.5 g/L. Plates were trypsinized at specific time points, and live cells were counted via trypan-blue exclusion using a BioRad TC20 Automated Cell Counter. Absolute cell counts were used to determine cell viability rather than the 3-(4,5-dimethylthiazol-2-yl)-2,5-diphenyltetrazolium (MTT) assay, as the latter uses mitochondrial metabolism as a marker of cell viability, and we sought to avoid any potential confounding effects of culture conditions on mitochondria.

### In vivo studies

GBM12, GBM164, and GBM196 cells were sourced from tumors propagated as subcutaneous growths in athymic nude mice and prepared for implantation. Briefly, after euthanizing the animal, tumors were aseptically excised from the flank and minced in a sterile culture dish with a scalpel. The cell suspension was centrifuged after mechanical disruption, filtered through 70 µM nylon-mesh filters, re-centrifuged, and re-suspended in an equal volume ratio of cell culture media to Matrigel. A 16-gauge needle and syringe were used to inject the cell suspension into the flank of NCr female athymic nude mice (NCRNU-F sp/sp, Taconic).

Intracranial injections of NPA and NPAC1 cells were performed as described previously in 10 week-old female C57BL/6J mice (Jackson) [14].

Animals received post-operative support care through administration of 0.9% saline solution, DietGel 76A, and thermal support while recovering from anesthesia until awake and ambulatory. One administration of meloxicam 1 mg/kg was given at the time of the procedure for 24 hour analgesia coverage, followed by another dose approximately 24 hours post-procedure, and a final dose 48 hours post-procedure if deemed necessary.

### Animal monitoring and treatment

All animals were fed standard rodent chow (standard diet, or SD) for 3 days following engraftment before being randomized to either remain on SD *ad libitum*, or given a formulated ketogenic diet (KD) (Ketogenic Diet TD.96355, Envigo Teklad Diets, Madison WI) *ad libitum*. The KD was received directly from the manufacturer and was a nutritionally complete diet composed of a 4.25 ratio of fat to protein plus carbohydrate (15.3% protein, 0.5% carbohydrate, 67.4% fat). Animals were placed on a weekly cyclic diet interchanging between SD and KD to control their weights and more effectively maintain plasma ketone levels, as described elsewhere [15]. Animals were monitored bi-weekly for signs of morbidity such as weight loss, behavioral changes, and hunched positions, and were euthanized when tumor size approached the recommended limit of 2000 mm^3^, or when moribund, via CO_2_ asphyxiation followed by cervical dislocation.

Flank tumor volume was based on caliper measurements, and calculated using the modified ellipsoidal formula [16, 17]: V=(length x width^2^)/2.

### Statistical Analyses

Differences between mean values of two groups were compared using two-sample *t*-test, or between multiple groups by one-way analysis of variance (ANOVA) and post hoc Tukey’s test; *P* values less than 0.05 were considered significant. Log-rank tests compared survival between groups. Graph generation and statistical analyses were performed with GraphPad Prism 5 (version 5.02, GraphPad Software, San Diego, CA).

## RESULTS

First, we examined the effects of a ketogenic-like diet on cultured IDH1^wt^ and IDH1^mut^ patient-derived cancer cell lines under the following conditions: (i) normal basal glucose (NG) of 4.5 g/L; (ii) low glucose (LG) of 1.0 g/L; (iii) NG with 10 mM β-hydroxybutyrate (BHB); (iv) LG with BHB (**Fig 1**). Neither IDH1^wt^ GBM6 nor IDH1^mut^ HT1080 cells showed any difference in cell number across the four culture conditions over time (**Fig 1A** and **Fig 1D**). Interestingly, IDH1^wt^ GBM43 and IDH1^mut^ TB09 cells showed the greatest cell number when grown in LG +BHB (**Fig 1B** and **Fig 1C**).

**Fig 1.**
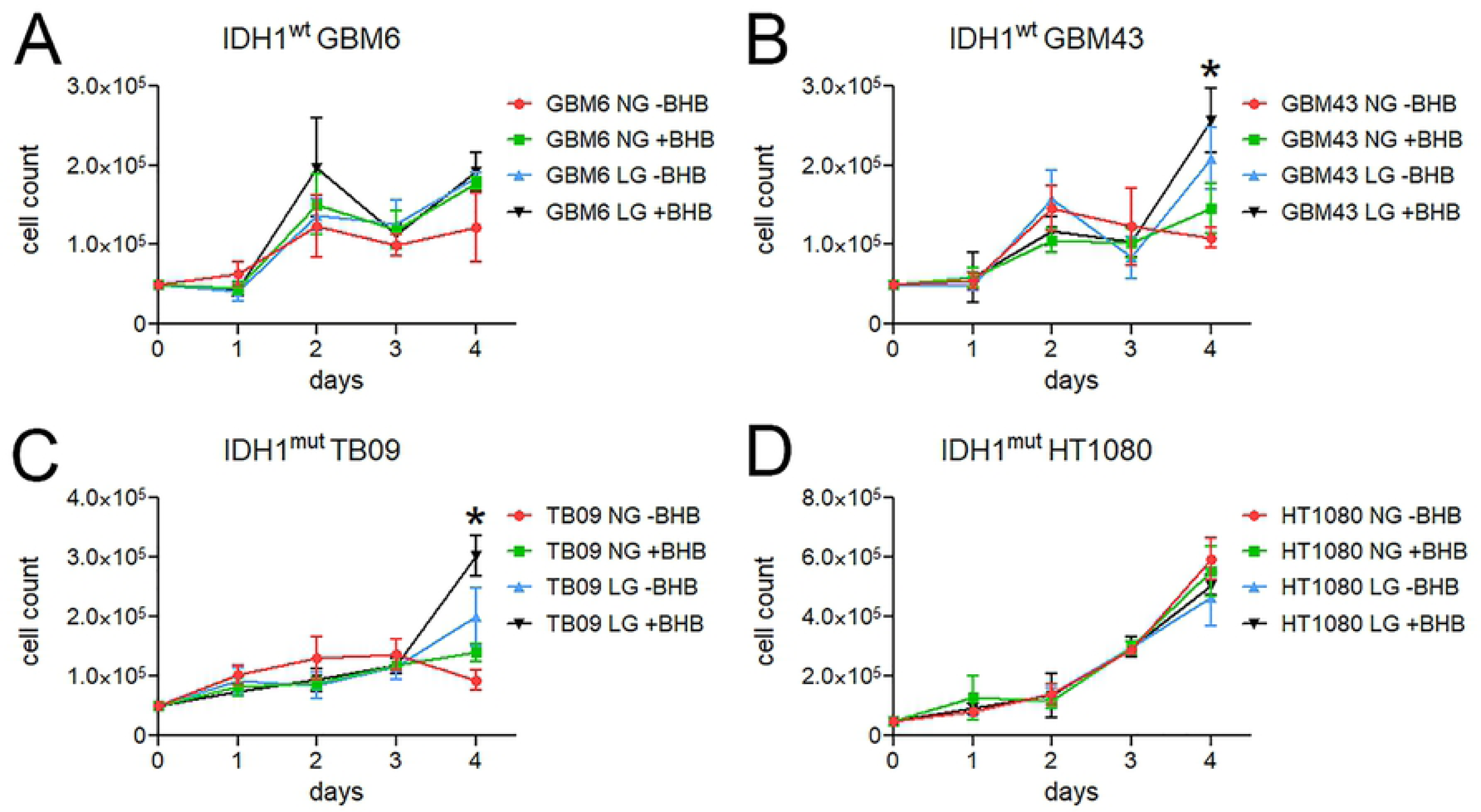
Effect of ketones on patient-derived IDH1^wt^ and IDH1^mut^ tumor cell proliferation *in vitro*. Absolute cell counts over four days of culture in either normal glucose (NG) or low glucose (LG), with or without 10 mM β-hydroxybutyrate (BHB). \**P*<0.05 by one-way ANOVA.

Next, we evaluated the ability of KD to affect the growth of patient-derived IDH1^wt^ and IDH1^mut^ GBM cells in the flanks of immunocompromised mice (**Fig 2**). (The flank was chosen because IDH1^mut^ GBM164 and GBM196 cells grow poorly as intracranial xenografts.) No significant difference in tumor volume was found in KD mice versus control among any of the tumor types.

**Fig 2.**
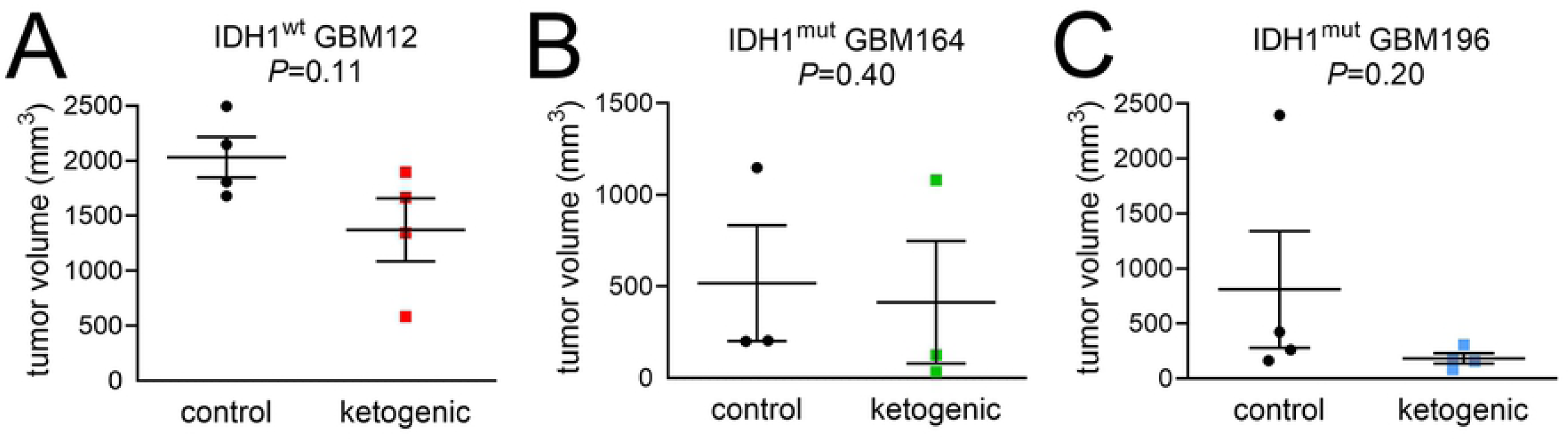
The effect of KD on patient-derived IDH1^wt^ and IDH1^mut^ flank xenograft growth. Tumor volume measurements for (A) IDH1^wt^ GBM12 at 33 days post-engraftment, (B) IDH1^mut^ GBM164 at 69 days, (C) IDH1^mut^ GBM 196 at 52 days, either on normal control diet or KD. Endpoints differed due to differing rates of tumor growth.

Finally, to assess the effect of KD on orthotopic gliomas in the presence of an intact immune system, we engrafted IDH1^wt^ NPA and IDH1^mut^ NPAC1 cells into the brains of immunocompetent mice (**Fig 3**). Among mice maintained on a regular diet, those engrafted with NPAC1 cells survived 20% longer than mice engrafted with NPA cells (median survival 26.5 days versus 22.0 days, HR=0.11, 95% CI=0.03-0.39, *P*=0.0022). However, KD had no effect on survival in either NPA-engrafted subjects (HR=0.47, 95% CI=0.15-1.5, *P*=0.27) (**Fig 3A**) or NPAC1-engrafted subjects (HR=1.0, 95% CI=0.31-3.2, *P*=0.81) (**Fig 3B**).

**Fig 3.**
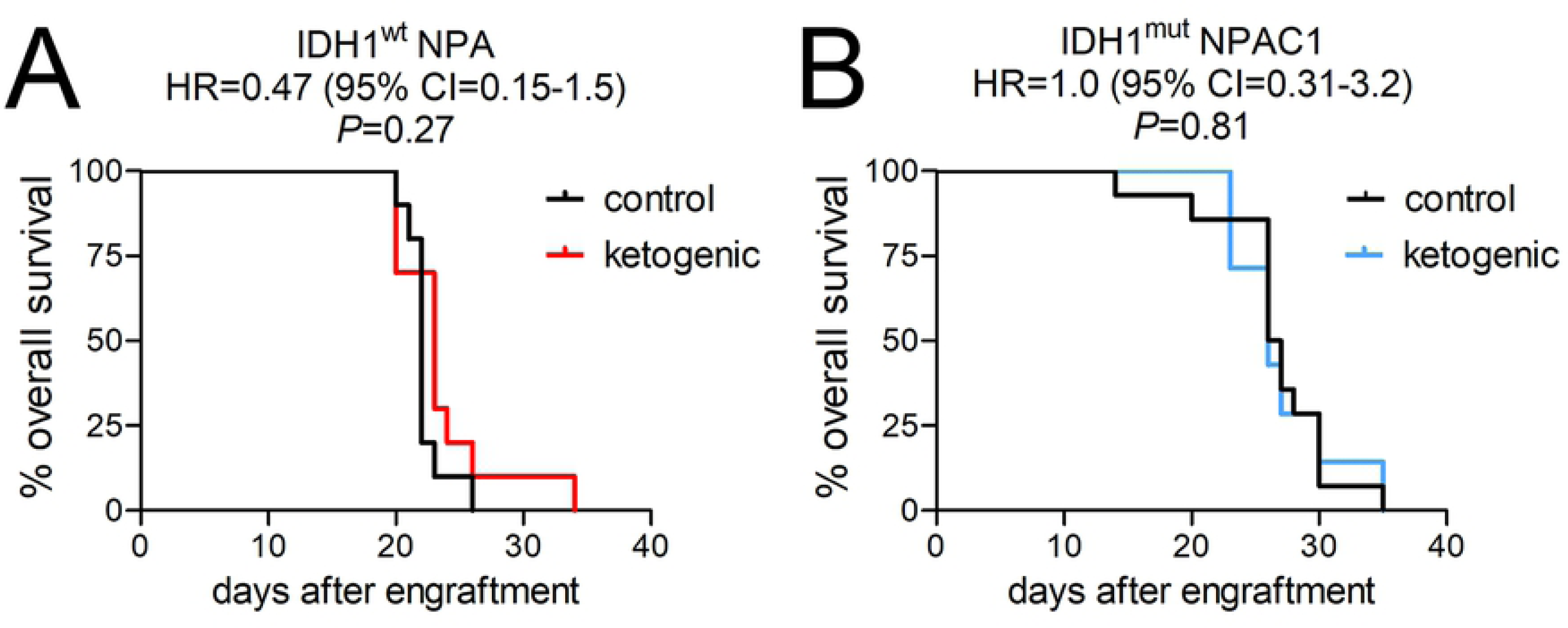
The effect of KD on patient-derived IDH1^wt^ and IDH1^mut^ intracranial xenograft growth. Kaplan-Meier survival curves of mice engrafted with (A) IDH1^wt^ NPA or (B) IDH1^mut^ NPAC1, either on normal control diet or KD.

## DISCUSSION

Altered metabolism in cancer cells raises the possibility of exploiting metabolic vulnerabilities to retard tumor growth. KD is designed to do this by depriving cancer cells of glucose, since normal cells are better able to utilize ketone bodies for their energy needs. While KD may indeed have efficacy against some cancers, clinical studies have been limited in gliomas [18]. One recurring limitation has been the lack of comparison between IDH1^wt^ and IDH1^mut^ glioma subtypes. A prior study by others suggested a potential sensitivity of IDH1^mut^ cancer to KD, based on *in vitro* data using engineered HCT116 cells [13]. Thus, we sought to determine KD efficacy in a variety of preclinical *in vitro* and *in vivo* models of IDH1^wt^ and IDH1^mut^ glioma. Our data suggest that KD does not have sufficient activity against either IDH1^wt^ or IDH1^mut^ glioma to warrant further investigation in clinical trials.

Since the original studies describing IDH1^mut^ and its neoenzymatic activity in cancer, a great deal of research has been done studying the metabolic effects of IDH1^mut^, often generating conflicting results. For example, whereas some have shown that IDH1^mut^ depletes the cell of TCA intermediates [19-24], others have found little to no change [10, 25, 26], and that IDH1^mut^ gliomas may use lactate and glutamate anaplerosis to replenish TCA intermediates [27]. Some have suggested that glycolysis is reduced in IDH1^mut^ gliomas [21, 25]. However, one group found reduced glucose uptake by IDH1^mut^ glioma cells, but otherwise no difference in the rate of glycolysis compared to IDH1^wt^ cells [26].

Results in IDH1^mut^ metabolic research seem to vary greatly depending on whether one is studying cells overexpressing IDH1^mut^ via transduction or is focusing on cells and tissues with endogenous, IDH1^mut^, as more pronounced metabolic changes tend to occur in the former than the latter [26]. This suggests that cells with endogenous IDH1^mut^ can, over time, adjust their metabolism to at least partially compensate for perturbations caused by the mutant enzyme. For example, IDH1^mut^ gliomas may compensate for the depletion of TCA intermediates by upregulating glutamate dehydrogenase 2 expression [21]. Since IDH1^mut^ gliomas mostly use the TCA precursor glutamine to produce D2HG, these tumors compensate by turning pyruvate into TCA intermediates [28, 29]. While IDH1^mut^ consumes NAPDH, which should lead to glutathione depletion, IDH1^mut^ gliomas upregulate enzymes involved in glutathione synthesis, thereby maintaining glutathione levels [30]. Thus, when IDH1^wt^ wild-type cells are abruptly forced to overexpress IDH1^mut^, any metabolic results, including sensitivity to KD-like conditions, need to be validated in patient-derived and/or transgenic models with endogenous IDH1^mut^.

Prolonged adherence to KD is notoriously difficult [31]. For a large well-controlled KD clinical trial to be justified in glioma patients, the preclinical evidence would therefore need to be particularly compelling. Our results suggest that KD is not a promising therapeutic strategy in glioma patients, regardless of IDH1^mut^ status.

## ACKNOWLEGMENTS

This work was supported by R01NS102669 and R01NS117104, the Northwestern University P50CA221747 SPORE in Brain Tumor Research, and the Lou and Jean Malnati Brain Tumor Institute.

## REFERENCES

1. Ostrom QT, Gittleman H, Liao P, Vecchione-Koval T, Wolinsky Y, Kruchko C, et al. CBTRUS Statistical Report: Primary brain and other central nervous system tumors diagnosed in the United States in 2010-2014. Neuro Oncol. 2017;19(suppl_5):v1-v88. Epub 2017/11/09. doi: 10.1093/neuonc/nox158. PubMed PMID: 29117289; PubMed Central PMCID: PMCPMC5693142.

2. Stupp R, Mason WP, van den Bent MJ, Weller M, Fisher B, Taphoorn MJ, et al. Radiotherapy plus concomitant and adjuvant temozolomide for glioblastoma. The New England journal of medicine. 2005;352(10):987-96. Epub 2005/03/11. doi: 10.1056/NEJMoa043330. PubMed PMID: 15758009.

3. Unruh D, Mirkov S, Wray B, Drumm M, Lamano J, Li YD, et al. Methylation-dependent Tissue Factor suppression contributes to the reduced malignancy of IDH1 mutant gliomas. Clin Cancer Res. 2018;28:1078–0432.

4. Burnet NG, Jefferies SJ, Benson RJ, Hunt DP, Treasure FP. Years of life lost (YLL) from cancer is an important measure of population burden--and should be considered when allocating research funds. Br J Cancer. 2005;92(2):241–5.

5. Hanahan D, Weinberg RA. Hallmarks of cancer: the next generation. Cell. 2011;144(5):646-74. PubMed PMID: 21376230.

6. Warburg O, Wind F, Negelein E. THE METABOLISM OF TUMORS IN THE BODY. J Gen Physiol. 1927;8(6):519-30. PubMed PMID: 19872213.

7. Parsons DW, Jones S, Zhang X, Lin JC, Leary RJ, Angenendt P, et al. An integrated genomic analysis of human glioblastoma multiforme. Science. 2008;321(5897):1807-12. PubMed PMID: 18772396.

8. Yan H, Parsons DW, Jin G, McLendon R, Rasheed BA, Yuan W, et al. IDH1 and IDH2 mutations in gliomas. The New England journal of medicine. 2009;360(8):765-73. PubMed PMID: 19228619.

9. Horbinski C. What do we know about IDH1/2 mutations so far, and how do we use it? Acta neuropathologica. 2013;125(5):621-36. Epub 2013/03/21. doi: 10.1007/s00401-013-1106-9 [doi]. PubMed PMID: 23512379.

10. Dang L, White DW, Gross S, Bennett BD, Bittinger MA, Driggers EM, et al. Cancer-associated IDH1 mutations produce 2-hydroxyglutarate. Nature. 2009;462(7274):739-44. PubMed PMID: 19935646.

11. Gano LB, Patel M, Rho JM. Ketogenic diets, mitochondria, and neurological diseases. J Lipid Res. 2014;55(11):2211-28. Epub 2014/05/23. doi: 10.1194/jlr.R048975. PubMed PMID: 24847102; PubMed Central PMCID: PMCPMC4617125.

12. Chung HY, Park YK. Rationale, Feasibility and Acceptability of Ketogenic Diet for Cancer Treatment. J Cancer Prev. 2017;22(3):127-34. Epub 2017/10/12. doi: 10.15430/jcp.2017.22.3.127. PubMed PMID: 29018777; PubMed Central PMCID: PMCPMC5624453.

13. Fu X, Chin RM, Vergnes L, Hwang H, Deng G, Xing Y, et al. 2-Hydroxyglutarate Inhibits ATP Synthase and mTOR Signaling. Cell Metab. 2015;22(3):508-15. Epub 2015/07/21. doi: 10.1016/j.cmet.2015.06.009. PubMed PMID: 26190651; PubMed Central PMCID: PMCPMC4663076.

14. Nunez FJ, Mendez FM, Kadiyala P, Alghamri MS, Savelieff MG, Garcia-Fabiani MB, et al. IDH1-R132H acts as a tumor suppressor in glioma via epigenetic up-regulation of the DNA damage response. Sci Transl Med. 2019;11(479). Epub 2019/02/15. doi: 10.1126/scitranslmed.aaq1427. PubMed PMID: 30760578; PubMed Central PMCID: PMCPMC6400220.

15. Newman JC, Covarrubias AJ, Zhao M, Yu X, Gut P, Ng CP, et al. Ketogenic Diet Reduces Midlife Mortality and Improves Memory in Aging Mice. Cell Metab. 2017;26(3):547-57 e8. PubMed PMID: 28877458.

16. Jensen MM, Jørgensen JT, Binderup T, Kjaer A. Tumor volume in subcutaneous mouse xenografts measured by microCT is more accurate and reproducible than determined by 18F-FDG-microPET or external caliper. BMC Med Imaging. 2008;8:16. Epub 2008/10/18. doi: 10.1186/1471-2342-8-16. PubMed PMID: 18925932; PubMed Central PMCID: PMCPMC2575188.

17. Faustino-Rocha A, Oliveira PA, Pinho-Oliveira J, Teixeira-Guedes C, Soares-Maia R, da Costa RG, et al. Estimation of rat mammary tumor volume using caliper and ultrasonography measurements. Lab Anim (NY). 2013;42(6):217-24. Epub 2013/05/22. doi: 10.1038/laban.254. PubMed PMID: 23689461.

18. Klement RJ, Brehm N, Sweeney RA. Ketogenic diets in medical oncology: a systematic review with focus on clinical outcomes. Medical oncology (Northwood, London, England). 2020;37(2):14. Epub 2020/01/14. doi: 10.1007/s12032-020-1337-2. PubMed PMID: 31927631.

19. Biedermann J, Preussler M, Conde M, Peitzsch M, Richter S, Wiedemuth R, et al. Mutant IDH1 Differently Affects Redox State and Metabolism in Glial Cells of Normal and Tumor Origin. Cancers. 2019;11(12). Epub 2020/01/01. doi: 10.3390/cancers11122028. PubMed PMID: 31888244; PubMed Central PMCID: PMCPMC6966450.

20. Miyata S, Tominaga K, Sakashita E, Urabe M, Onuki Y, Gomi A, et al. Comprehensive Metabolomic Analysis of IDH1(R132H) Clinical Glioma Samples Reveals Suppression of β-oxidation Due to Carnitine Deficiency. Sci Rep. 2019;9(1):9787. Epub 2019/07/07. doi: 10.1038/s41598-019-46217-5. PubMed PMID: 31278288; PubMed Central PMCID: PMCPMC6611790.

21. Waitkus MS, Pirozzi CJ, Moure CJ, Diplas BH, Hansen LJ, Carpenter AB, et al. Adaptive Evolution of the GDH2 Allosteric Domain Promotes Gliomagenesis by Resolving IDH1(R132H)-Induced Metabolic Liabilities. Cancer Res. 2018;78(1):36-50. Epub 2017/11/04. doi: 10.1158/0008-5472.Can-17-1352. PubMed PMID: 29097607; PubMed Central PMCID: PMCPMC5754242.

22. Ohka F, Ito M, Ranjit M, Senga T, Motomura A, Motomura K, et al. Quantitative metabolome analysis profiles activation of glutaminolysis in glioma with IDH1 mutation. Tumour Biol. 2014;35(6):5911-20. Epub 2014/03/05. doi: 10.1007/s13277-014-1784-5. PubMed PMID: 24590270.

23. Reitman ZJ, Jin G, Karoly ED, Spasojevic I, Yang J, Kinzler KW, et al. Profiling the effects of isocitrate dehydrogenase 1 and 2 mutations on the cellular metabolome. Proc Natl Acad Sci U S A. 2011;108(8):3270-5. PubMed PMID: 21289278.

24. Grassian AR, Parker SJ, Davidson SM, Divakaruni AS, Green CR, Zhang X, et al. IDH1 mutations alter citric acid cycle metabolism and increase dependence on oxidative mitochondrial metabolism. Cancer Res. 2014;74(12):3317-31. Epub 2014/04/24. doi: 10.1158/0008-5472.Can-14-0772-t. PubMed PMID: 24755473; PubMed Central PMCID: PMCPMC4885639.

25. Zhou L, Wang Z, Hu C, Zhang C, Kovatcheva-Datchary P, Yu D, et al. Integrated Metabolomics and Lipidomics Analyses Reveal Metabolic Reprogramming in Human Glioma with IDH1 Mutation. J Proteome Res. 2019;18(3):960-9. Epub 2019/01/01. doi: 10.1021/acs.jproteome.8b00663. PubMed PMID: 30596429.

26. Garrett M, Sperry J, Braas D, Yan W, Le TM, Mottahedeh J, et al. Metabolic characterization of isocitrate dehydrogenase (IDH) mutant and IDH wildtype gliomaspheres uncovers cell type-specific vulnerabilities. Cancer Metab. 2018;6:4. Epub 2018/04/26. doi: 10.1186/s40170-018-0177-4. PubMed PMID: 29692895; PubMed Central PMCID: PMCPMC5905129.

27. Khurshed M, Molenaar RJ, Lenting K, Leenders WP, van Noorden CJF. In silico gene expression analysis reveals glycolysis and acetate anaplerosis in IDH1 wild-type glioma and lactate and glutamate anaplerosis in IDH1-mutated glioma. Oncotarget. 2017;8(30):49165-77. Epub 2017/05/04. doi: 10.18632/oncotarget.17106. PubMed PMID: 28467784; PubMed Central PMCID: PMCPMC5564758.

28. Izquierdo-Garcia JL, Viswanath P, Eriksson P, Cai L, Radoul M, Chaumeil MM, et al. IDH1 Mutation Induces Reprogramming of Pyruvate Metabolism. Cancer Res. 2015;75(15):2999-3009. Epub 2015/06/06. doi: 10.1158/0008-5472.Can-15-0840. PubMed PMID: 26045167; PubMed Central PMCID: PMCPMC4526330.

29. Izquierdo-Garcia JL, Cai LM, Chaumeil MM, Eriksson P, Robinson AE, Pieper RO, et al. Glioma cells with the IDH1 mutation modulate metabolic fractional flux through pyruvate carboxylase. PLoS One. 2014;9(9):e108289. Epub 2014/09/23. doi: 10.1371/journal.pone.0108289. PubMed PMID: 25243911; PubMed Central PMCID: PMCPMC4171511.

30. Fack F, Tardito S, Hochart G, Oudin A, Zheng L, Fritah S, et al. Altered metabolic landscape in IDH-mutant gliomas affects phospholipid, energy, and oxidative stress pathways. EMBO Mol Med. 2017;9(12):1681-95. Epub 2017/10/22. doi: 10.15252/emmm.201707729. PubMed PMID: 29054837; PubMed Central PMCID: PMCPMC5709746.

31. Brouns F. Overweight and diabetes prevention: is a low-carbohydrate-high-fat diet recommendable? Eur J Nutr. 2018;57(4):1301-12. PubMed PMID: 29541907.

